# Shared regulatory pathways reveal novel genetic correlations between grip strength and neuromuscular disorders

**DOI:** 10.1101/765495

**Authors:** Sreemol Gokuladhas, William Schierding, David Cameron-Smith, Melissa Wake, Emma L. Scotter, Justin O’Sullivan

## Abstract

**Background:** Muscle weakness and muscle wasting can be a consequence of aging (sarcopenia) and neuromuscular disorders (NMD). Genome-wide association (GWA) studies have identified genetic variants associated with grip strength (GS, an inverse measure of muscle weakness) and NMD (multiple sclerosis (MS), myasthenia gravis (MG) and amyotrophic lateral sclerosis (ALS)). However, how these variants contribute to the muscle weakness caused by aging or NMD remains obscure.

**Methods:** We have integrated GS and NMD associated SNPs in a multimorbid analysis that leverages high-throughput chromatin interaction (Hi-C) data and expression quantitative trait loci (eQTL) data to identify allele-specific gene regulation (*i.e.* eGenes). Pathways and shared drug targets that are enriched by colocalised eGenes were then identified using pathway and drug enrichment analysis.

**Results:** We identified gene regulatory mechanisms (eQTL-eGene effects) associated with GS, MG, MS and ALS. The eQTLs associated with GS regulate a subset of eGenes that are also regulated by the eQTLs of MS, MG and ALS. Yet, we did not find any eGenes commonly regulated by all four phenotypes associated eQTLs. By contrast, we identified three pathways (mTOR signaling, axon guidance, and alcoholism) that are commonly affected by the gene regulatory mechanisms associated with all four phenotypes. 13% of the eGenes we identified were known drug targets, and GS shares at least one druggable eGene and pathway with each of the NMD phenotypes.

**Conclusions:** Collectively, these findings identify significant biological overlaps between GS and NMD, demonstrating the potential for spatial genetic analysis to identify mechanisms underlying muscle weakness due to aging and NMD.

## Introduction

Age-associated changes in skeletal muscle structure and composition are influenced by various factors including genetics, physical inactivity, malnutrition, hormonal and environmental changes[1–3]. Understanding the biological pathways that contribute to disease and age-associated muscle wasting is critical if we are to develop effective therapies. Grip strength (GS) is strongly correlated with age-related loss of muscle strength (sarcopenia) and therefore is used as a predictor of health-related quality of life[4,5]. Cachexia is a primary characteristic in a number of chronic neuromuscular diseases (NMD), including myasthenia gravis (MG), amyotrophic lateral sclerosis (ALS), and multiple sclerosis (MS)[6–8]. For all of these diseases, the primary disease diagnosis is complicated by reduced physical function, reduced tolerance for therapy, increased burden on the healthcare system, and increased mortality that comes with cachexia[9].

Numerous mechanisms have been suggested to be involved in NMD-related muscle weakness, including mitochondrial dysfunction[10], reduced numbers of motor units[11], and functional deficits at the neuromuscular junction (NMJ)[12]. In MG, auto-antibodies are aberrantly directed against acetylcholine receptors, muscle-specific kinase and lipoprotein-related protein 4 in the postsynaptic membrane at the NMJ, disrupting cholinergic signal transmission, thereby inducing muscle weakness[13,14]. By contrast, ALS is a fatal motor neuron disease that causes progressive degeneration of upper and lower motor neurons in the brain and spinal cord leading to muscle weakness, paralysis and death. Several lines of evidence suggest that there are also autonomous changes in skeletal muscle in ALS, independent of motor neuron degeneration. In ALS, skeletal muscle harbours the characteristic protein aggregates (composed principally of TDP-43)[15] whose burden in the CNS predicts neurodegeneration[16,17]. In muscle, these TDP-43 protein aggregates may disrupt the transport of mRNAs essential for muscle fibre regeneration[18,19]. Together these findings support a direct contribution of skeletal muscle to the ALS phenotype. Similarly, changes in the characteristics of skeletal muscle are observed in MS, an autoimmune-mediated demyelinating disease of the central nervous system that leads to muscle disuse and weakness. Altered muscle function in MS includes impaired excitation-contraction coupling[20], reduced oxidative capacity of the skeletal muscle[21], slowing of muscle contraction[22], and decreased cross-sectional area of type I and type II muscle fibres[23]. Co-occurrences of MG:MS[24,25], MG:ALS[26–28] and MS:ALS[29,30] in patients and families have been reported, consistent with an etiological relationship or multimorbidity between these diseases. We hypothesize that GS, MG, MS and ALS share common biological pathways consistent with their common age-associated and/or disease-associated muscle wasting and weakness.

To date, there has been only one GWA study on the genetic features of cachexia, and only for body mass loss in COPD and cancer[31]. However, GWA studies have identified genetic variations (*i.e.* single nucleotide polymorphisms (SNPs)) associated with GS[32], MG[33], MS[34] and ALS[35]. These studies employed case-control approaches to identify the SNPs that are associated with these disorders. Most of these SNPs are located in the non-protein coding or regulatory regions of the genome[36]. The rarity of SNPs within coding sequences increases the difficulty of understanding how they function to impact on phenotype. One putative function is through alterations in regulatory regions (*e.g*. enhancers) through alteration of allele-specific long-range DNA contacts. Thus, the regulatory regions can be separated from their target genes by great distances [37]. Integrating the three-dimensional organisation of the genome, captured by proximity ligation (*e.g*. Hi-C) [38], with functional data (*e.g*. eQTLs) can identify the targets of these long-range regulatory interactions[39.

We hypothesize that SNP overlap between GS, MG, MS and ALS is uncommon, but that intersecting gene regulation and biological pathways explain the common features of the diseases (i.e. NMD and muscle wasting). In the present study, we identify the genes that are regulated by GS-, MG-, MS- and ALS-associated SNPs using high throughput chromatin interaction data (Hi-C) from 8 immortalized cell-lines and psoas muscle tissue (*i.e.* representative of the genomic contacts seen across cell types or only in skeletal muscle, respectively). Combining these spatial interaction data with functional eQTL data, we identify gene regulatory (*i.e.* eQTL-eGene) interactions associated with each phenotype. Subsequent pathway analyses identified the biological pathways associated with these phenotypes. Collectively, our analysis identifies numerous genetic and biological pathway overlaps between neuromuscular diseases (MS, MG and ALS) and GS, including three pathways that overlap all four. This suggests common and distinct genetic mechanisms underlying muscle weakness associated with age and disease.

## Methods

### Identification of functionally significant spatial regulatory interactions

Overall, 179, 18, 286 and 135 SNPs were associated with GS, MG, MS and, ALS, respectively, at a suggestive level of significance (p ≤ 5×10^−6^). Using the CoDeS3D algorithm[40], we mapped spatial regulatory connections between each phenotype-associated SNP and one or more gene coding regions (GENCODE release 19). For each SNP-gene pair, spatial contacts were identified from existing Hi-C[38] chromatin contact data derived from eight immortalized cell lines representing human germ layer lineages (GM12878, IMR90, HMEC, NHEK, K562, HUVEC, HeLa and KBM7)[41] and one tissue (Psoas muscle)[42] **(Table S1)**. Regulatory activity was supported by expression Quantitative Trait Loci (eQTL) in the expression level of partner genes (eGenes) in the Genotype-Tissue Expression (GTE×)[43] database (Version 7) using a False Discovery Rate [FDR] < 0.05[40]. Chromosome positions of eQTLs and eGenes were annotated according to Human reference genome GRCh37/hg19 assembly.

### Annotation of significant eGenes

Functional annotation of the eGenes was performed using the Gene Ontology (GO) Consortium web-based knowledgebase[44,45] (http://geneontology.org/, February 02, 2019). Overrepresentation was performed using PANTHER and the reference set of all *Homo sapiens* protein-coding genes. P-values were subjected to Bonferroni multiple testing correction with p< 0.05 being deemed significant.

### Identification of enriched pathways

iPathwayGuide (https://advaitabio.com/ipathwayguide/) was used to identify biological pathways that were enriched with colocalized eGenes. Analyses were performed using default parameters as described by Ahsan and Draghici (FDR < 0.05)[46].

### Identification of overlapping eGenes across phenotypes

R studio (Version 1.2) was used to compare eGene lists for each phenotype to identify intersections between all possible phenotype combinations. To test if gene overlaps were non-random, we performed a bootstrap analysis using 10000 randomly chosen sets of genes (Gencode v19) of the same size as the number of eGenes for each phenotype.

### Retrieving information about druggability of eGenes

The druggable eGenes were identified by querying all eGenes against the Drug Gene Interaction database (DGIdb, http://www.dgidb.org/) which integrates information on gene druggability and drug-gene interactions from 30 different sources[47,48].

## Results

### SNPs associated with generalized muscle weakness (GS) and neuromuscular disorders (MG, MS and ALS) form an overlapping functional regulatory network

SNPs associated with GS were identified from the published literature and the details are provided in the (**Table S2**). MG, MS, and ALS associated SNPs were obtained from GWAS catalog (https://www.ebi.ac.uk/gwas, January 20, 2018) and were analyzed through CoDeS3D[40] pipeline. The pipeline uses chromatin interaction (Hi-C) data from immortalized cell lines[41] to connect GS-, MG-, MS- and ALS-associated SNPs with eQTL data (GTEx). Despite the common muscle weakness phenotype presented by these disorders, there were no common SNPs shared between the sets of disease associated SNPs for each of these disorders. We identified SNP-gene spatial connections associated with eQTLs for GS (n=179), MG (n=18), MS (n=285), and ALS (n=135) SNPs **(Figure 1a)**. Approximately 95% of the spatial eQTL-eGene regulatory interactions identified across each phenotype occur between proximal (cis; ≤ 1 Mb between the SNP and the gene) rather than distal (trans; > 1 Mb between the SNP and gene, or the SNP and gene are located on different chromosomes) regions **(Figure 1b, Table S3)**. We then tested for phenotype-specific enrichment in the proportion of eQTLs versus non-eQTLs. While nearly 80% of the SNPs associated with GS, MG and MS are eQTLs, less than 50% of ALS-associated SNPs are eQTLs **(Figure 1c).**

**Figure 1.**
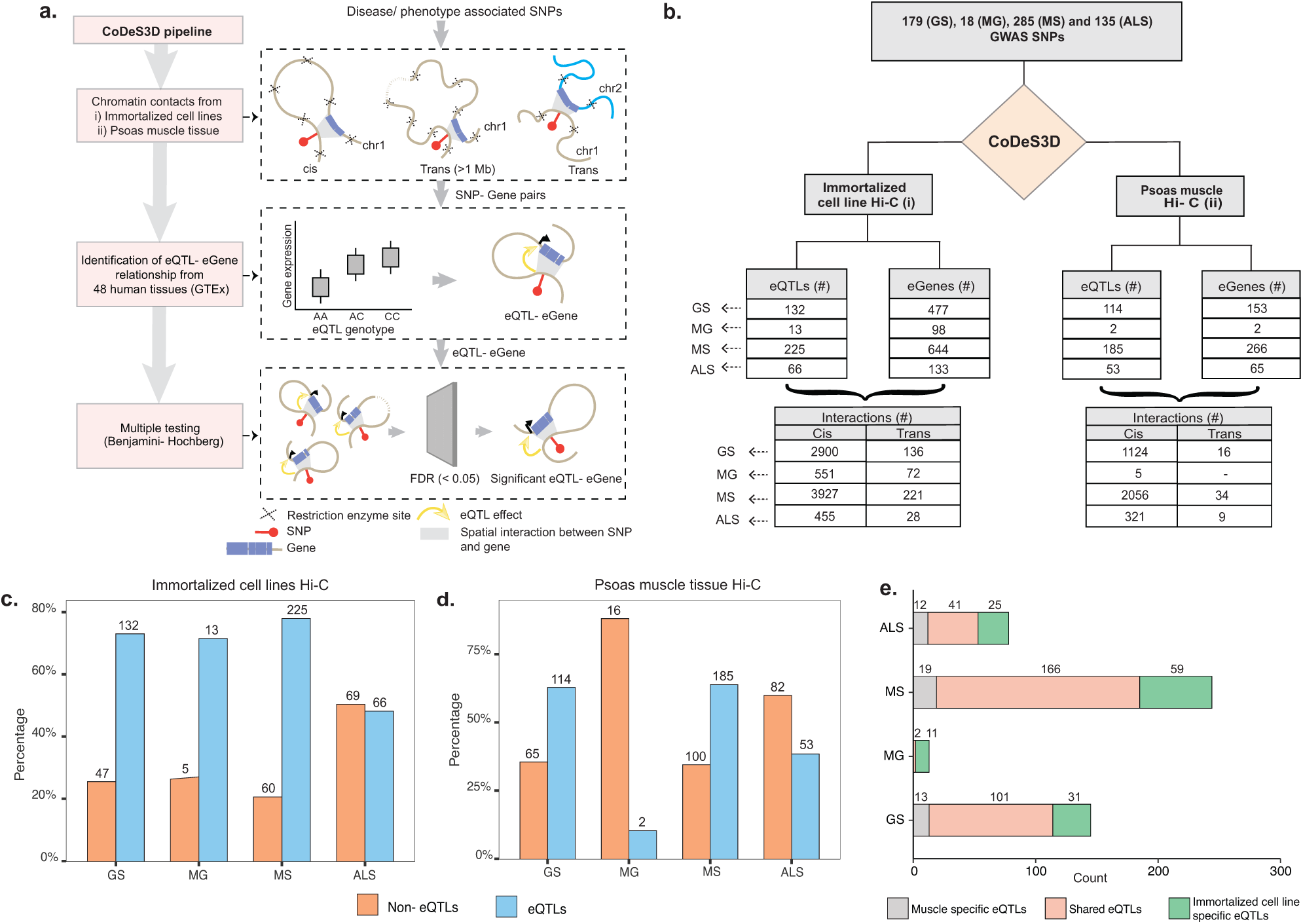
GWAS SNPs associated with GS, MG, MS, and ALS mark functional regulatory loci. **a.** CoDes3D workflow: The regulatory SNP-gene interactions (cis and trans) were identified from existing Hi-C datasets (*i.e.*, immortalized cell lines dataset from Rao *et al*[41] and psoas muscle tissue dataset from Schmitt *et al*[42]). The SNP-gene pairs were tested against the GTE× database to identify eQTL effects on the spatially connected genes. Those eQTL-eGenes with FDR < 0.05 were considered significant. **b**. Summary statistics of the regulatory interactions (spatial eQTLs) identified between phenotype (GS, MG, MS and ALS)-associated SNPs and eGenes. eQTLs were identified by integrating spatial information from Hi-C contact maps for immortalized cell lines (Rao et al[41]; left) or psoas muscle (Schmitt et al[42]; right) with eQTL data from GTEx (FDR < 0.05). Cis eQTLs are located ≤ 1 Mb from the target eGene. Trans eQTLs are located > 1 Mb from the target eGene or on a different chromosome. **c and d**. The proportion of eQTL: non-eQTL SNPs identified using **c.** the immortalized cell line Hi-C datasets[41], or **d**. the psoas muscle Hi-C dataset[42] **e.** Numbers of eQTLs specific to immortalized cell line and psoas muscle, or both Hi-C datasets

We hypothesized that the emergent genome organization in muscle would be more relevant to gene regulation in understanding the biological underpinnings of sarcopenia and NMD. Therefore, we repeated the CoDeS3D analysis using genomic organization data captured in psoas muscle tissue[42]**(Figure 1b (right); Table S4)**. We identified phenotype-specific enrichment of eQTLs in the psoas muscle Hi-C dataset **(Figure 1d)**. Furthermore, consistent with our hypothesis, and despite the Rao dataset being nearly 10 times larger, a considerable number of the identified regulatory interactions were specific to muscle **(Figure 1e, Figure S1).** Notably, we did not identify any eQTLs specific to psoas muscle using SNPs associated with MG. This suggests that MG-associated SNPs do not make a major contribution to the regulatory networks within skeletal muscle

### Shared eGenes between GS, MG, MS and ALS suggest common genetic mechanisms

The clinical manifestations, diagnoses, and treatments of MG, MS and ALS differ. However, despite having no overlapping SNPs, our analysis identified 89 shared eGenes between these phenotypes and generalized muscle weakness (GS**; Figure 2, Table S5)**. The degree of eGene overlap between phenotypes was dependent upon the Hi-C datasets that were used to identify the eQTLs (compare **Figures 2a** and **2b**). Bootstrapping (10000 test intersections) confirmed that the shared eGene overlaps identified using spatial SNP-gene sets captured within the immortalized cell lines were significantly greater than expected by chance **(Figure 2c).** Similarly, comparisons of eGenes identified using psoas muscle Hi-C data identified significant overlaps between the GS and MS, and MG and ALS SNP-gene sets **(Figure 2d)**.

**Figure 2.**
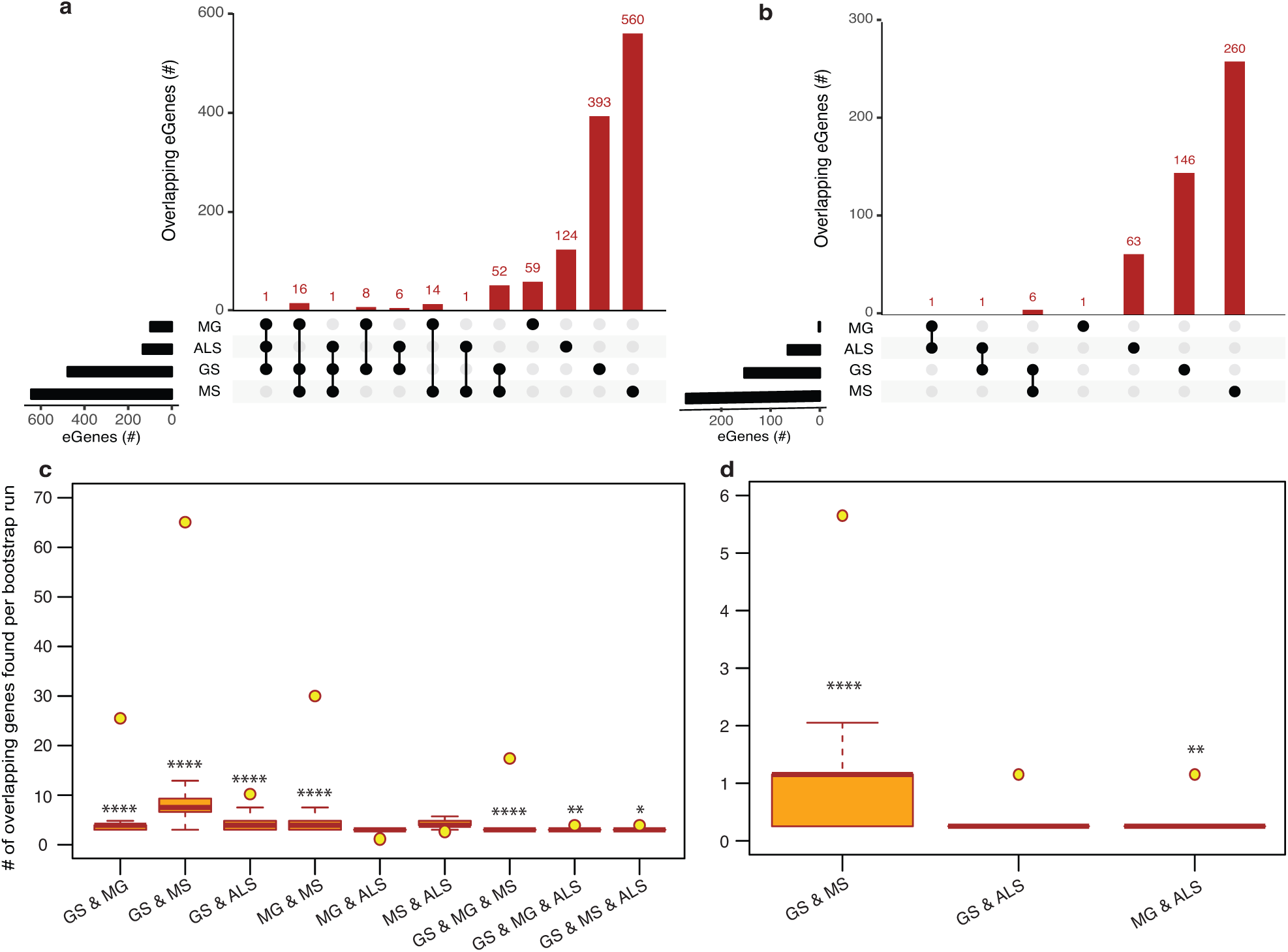
Comparisons of spatial regulatory interactions identified eGenes that are shared between GS, MG, MS and ALS. Overlap between eGene sets for GS, MG, MS and ALS identified using **a**. immortalized cell line Hi-C datasets, or **b.** the Psoas muscle Hi-C dataset. The total number of eGenes contained in each individual set is represented as horizontal bars (black). The dot matrix denotes phenotype overlap, where the line connecting dots indicates the sets being compared. **c**. and **d**. Boxplots showing the mean and range of bootstrapping values (n=10000) for overlaps between randomly selected gene sets to validate overlaps from **c.** the immortalized cell line Hi-C dataset, or **d.** the Psoas muscle Hi-C dataset. The yellow point represents the observed number of eGene overlaps between the stated phenotypes. **** p-value < 1×10^−5^ ** p-value < 0.01 *p-value <0.05

We identified 30 shared eGenes (mainly HLA loci genes) between MG and MS eQTLs; *ERMP1* and *SLC25A12* were regulated by MS and ALS eQTLs; and *C18orf8* was regulated by MG and ALS eQTLs **(Table S5)**. The 25 eGenes in common between GS and MG were overrepresented in immune system-related processes, including antigen processing and presentation, interferon-gamma-mediated signaling, T-cell receptor signaling and adaptive immune responses **(Table S6a)**. Similarly, ontological analysis of the 69 eGenes in common between GS and MS identified enrichment for immune system functions **(Table S6b)**. Collectively, these findings are consistent with a partially shared genetic aetiology underlying the development of muscle weakness between sarcopenia and NMD.

### Overlapping GS, MG, MS and ALS molecular pathways reveal putative therapeutic targets

Gradual muscle wasting and weakness is a comorbidity that we hypothesized would be reflected in shared biological pathways between MS, MG, ALS and GS. Using iPathwayGuide, we identified considerable overlap at the pathway levels between GS and NMD associated eGenes **(Figure 3)**. For example, GS (22 out of 42) and MG (16 out of 24) eGenes (immortalized cell line Hi-C dataset) are highly enriched for immune system and related pathways **(Figure S2, Table S7a).** Similarly, GS and MS eGenes are enriched for immune system, signal transduction, nervous system, and endocrine system-associated pathways **(Figure S3, Table S7b)**. eGenes for GS and ALS co-occurred within signal transduction, substance dependence, and cell motility pathways **(Figure S4, Table S7c)**.

**Figure 3.**
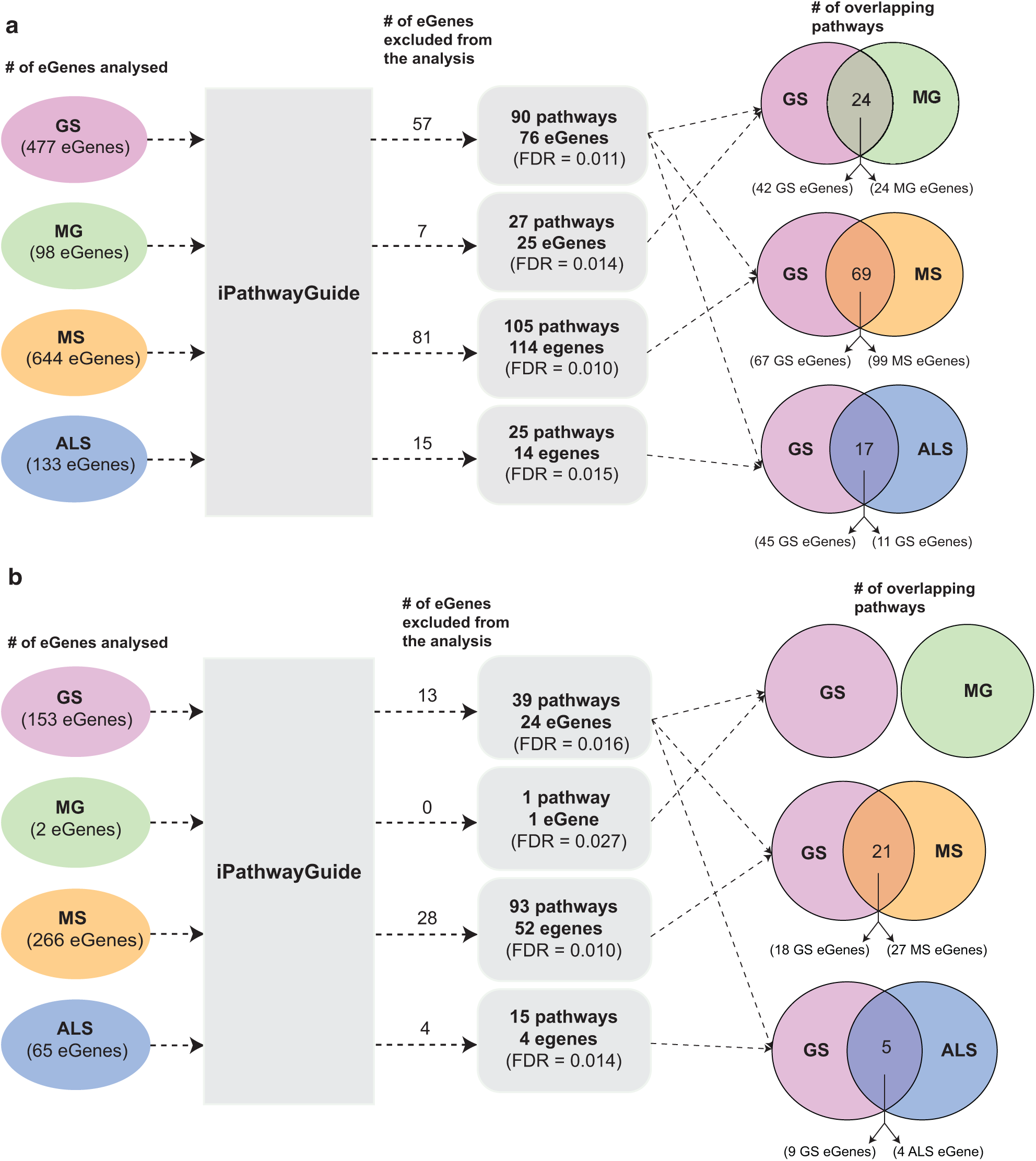
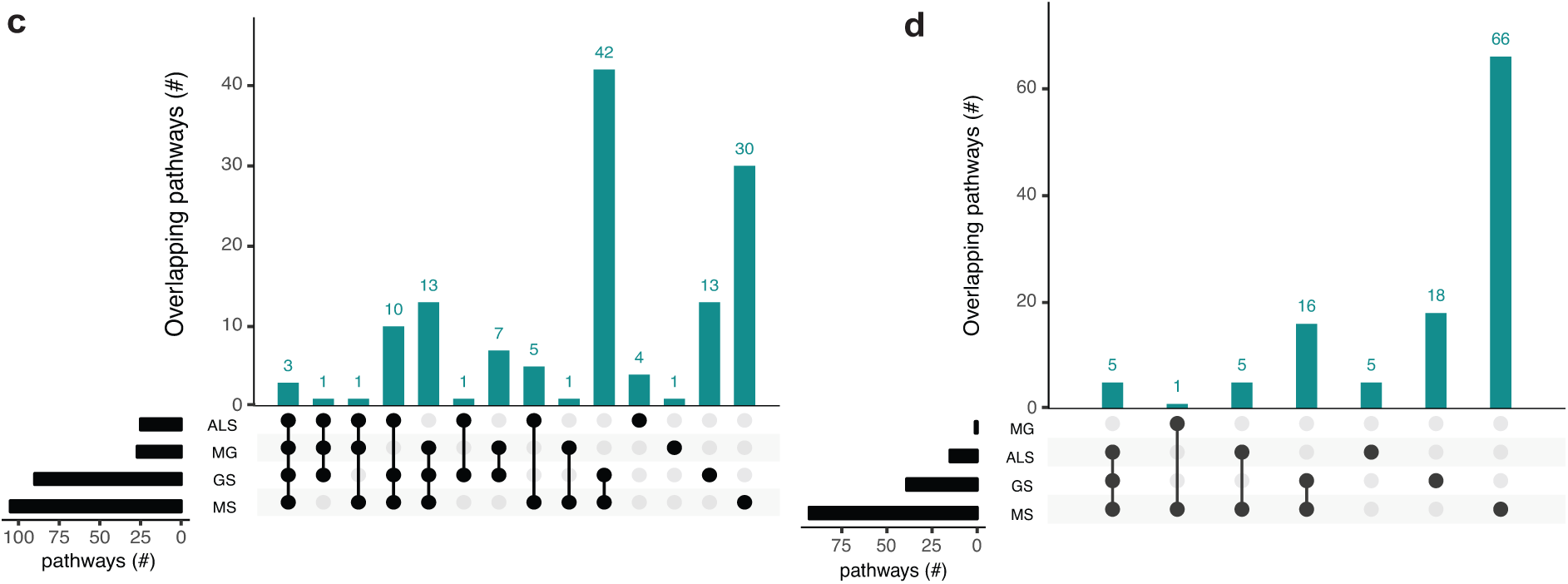
Co-occurrence of condition-specific eGenes within biological pathways is associated with comorbidity between GS, MG, MS and ALS. Overlap of biological pathways that contain phenotype-associated eGenes identified using chromatin interactions from **a.** immortalized cell lines, or **b.** the psoas muscle. Numbers of overlapping pathways enriched with eGenes from GS, MG, MS and ALS identified from **c.** immortalized cell line Hi-C dataset and **d.** psoas muscle tissue Hi-C dataset are plotted on the y axis. The total number of pathways impacted by each set of eGenes is represented as horizontal bars (black).

Shared eGenes explained some of the observed co-occurrence in pathways; > 50% of the shared pathways observed for GS:MG and GS:MS eGenes (immortalized cell line Hi-C datasets) contained both shared and condition-specific eGenes. By contrast, 99% of shared pathways identified for GS and ALS were due to co-occurrence of condition-specific eGenes. Similarly, all of the overlapping pathways (GS:MS and GS:ALS) identified using eGenes that were derived from the psoas muscle dataset were due to the co-occurrence of condition-specific eGenes (**Table S7d and S7e)**. Thus, pathway overlap was largely due to the co-occurrence of condition-specific eGenes within the same functional networks.

eGenes associated with NMD also co-occur in shared pathways, consistent with the existence of multimorbidity between these conditions **(Table S8)**. Notably, while there were no eGenes that were shared by all four phenotypes **(Figure 2a)**, there were three pathways (*i.e.* axon guidance, alcoholism and the mTOR signaling pathway) that contained co-occurring eGenes from all four conditions **(Figure 3c)**. For example, the axon guidance pathway contained eGenes that were regulated by genetic variants from one or multiple phenotypes **(Figure 4a).** Similar patterns were observed in both the mTOR signaling **(Figure 4b)** and alcoholism pathways **(Figure S5)**.

**Figure 4.**
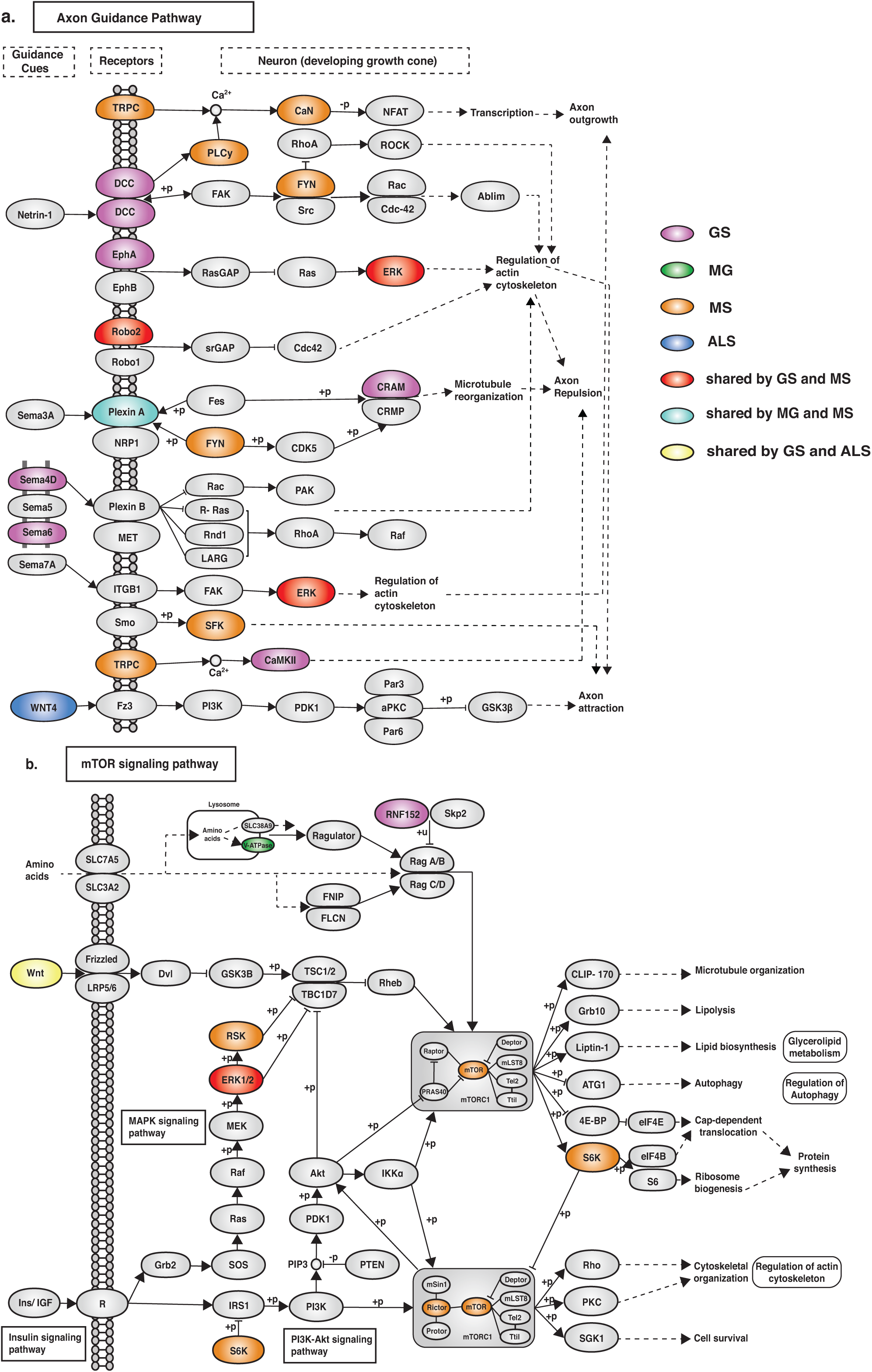
eGenes from all four phenotypes (GS, MG, MS and ALS) co-occurred within the axon guidance and mTOR signalling pathways. **a.** The axon guidance pathway is involved in neural development and defects are associated with neuronal disorders. **b.** The mTOR signaling pathway controls cellular processes including: cell growth, protein synthesis, cytoskeletal organisation, and autophagy[49]. GS-, MG-, MS- and ALS-associated eQTLs spatially regulate multiple eGenes that encode proteins involved in these pathways. Notably, proteins encoded by ALS eGenes are upstream regulators in both pathways.

The eGenes associated with GS, MG, MS and ALS that were identified using the psoas muscle dataset also co-occurred in biological pathways **(Figure 3d)**. The shared biological pathways that contained eGenes identified using the psoas muscle Hi-C dataset represented a subset of overlapping pathways identified using the eGenes from the immortalized cell line Hi-C dataset (**Table S8b)**.

### Overlapping, treatable biological pathways are shared drug targets for GS, MG, MS and ALS

eGenes that encode druggable proteins may have therapeutic value, especially as targets for drug repurposing or for the identification of potential targets for shared drug morbidity. Using the DGI database, we determined that 13% of the eGenes we identified are druggable **(Table S9 and S10)**. Notably, GS shares at least one druggable eGene and pathway with each of the NMD phenotypes **(Figure 5, Table S11a and S11b)**. While there were no druggable eGenes between ALS and MG, there was one druggable pathway (i.e. cytokine-cytokine receptor interaction pathway) that was shared between them **(Table S11c)**.

**Figure 5.**
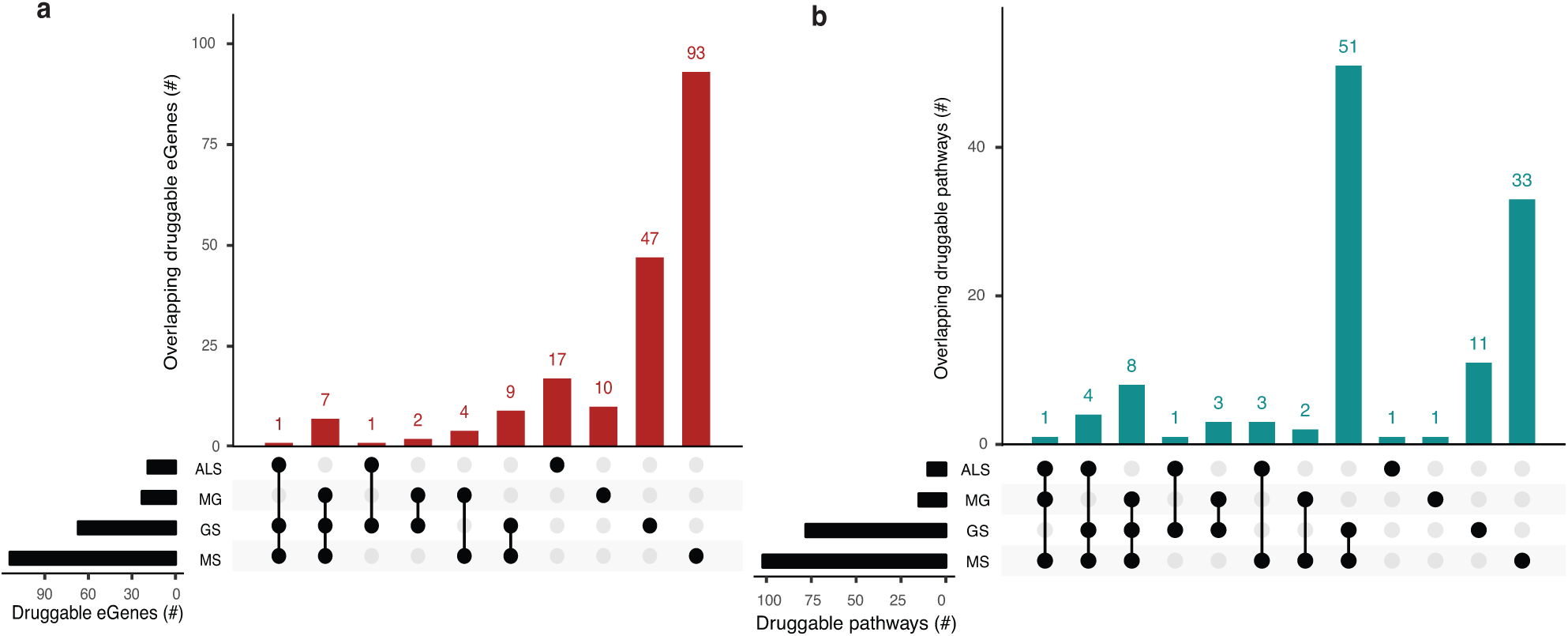
Phenotypes share druggable eGenes and druggable pathways. Overlapping **a.** druggable eGenes and **b.** pathways they occur in, as identified from the immortalized cell line Hi-C dataset. The number of druggable eGenes identified for each phenotype (left) and the number of druggable pathways identified for each phenotype (right) are represented as horizontal bars. The matrix denotes the intersection between sets under comparison.

We identified entirely unique sets of druggable eGenes for GS (4 eGenes), MS (18 eGenes) and ALS (1 eGene) from the psoas muscle that were not found in immortalized cell lines **(Table S9)**. Three out of four GS druggable eGenes co-occur within the neuroactive ligand-receptor interaction pathway. Nine out of eighteen MS druggable eGenes were involved in 25 different pathways, predominantly in signal transduction pathways **(Table S11d)**. From the psoas muscle Hi-C dataset we identified unique druggable eGenes (e.g. GRID1, TNFRSF1A, STAT4) which represent potential therapeutic targets for treating the muscle wasting/weakness symptoms associated with NMD and sarcopenia.

## Discussion

This is the first report of shared biological mechanisms between genetic variants associated with the generalized age-associated decline of muscle strength (as measured by hand GS) and muscle weakness/wasting caused by neuromuscular disorders (i.e. MG, MS and ALS). The existence of shared eGenes, whose expression is correlated with condition-associated SNPs, is consistent with subtle changes within combinatorial gene regulatory mechanisms contributing to the risk of generalized aspects of muscle weakness. Moreover, the observation of the co-occurrence of condition-specific eGenes within shared biological pathways provides insights into the mechanisms associated with the risk of development of generalized age- and disease-associated muscle weakness.

### Shared immune markers highlight pathways of muscle protein synthesis

Altered concentrations of circulating immune markers and heightened immune responses have been shown to lead to reduced muscle protein synthesis[50]. We identified eGenes that contribute to immune system-related functions and whose expression levels were correlated with genetic variants associated with GS and MG or MS. The co-occurrence of eGenes (e.g. HLA loci genes, *C2*, *TAP2*, *MICB*) specific to each of the phenotypes within immune system-related pathways is consistent with these pathways having a significant role in the development of age and disease-related muscle weakness. Similarly, we determined that the expression level of the *SLC25A12* gene is associated with rs2044469 (GS), rs4953911 and rs882300 (MS) and, rs1400816 (ALS). Notably, alterations to *SLC25A12* expression affect mitochondrial structure, function and biogenesis and are associated with age-related muscle dysfunction[51]. Collectively, these observations highlight the complex interrelationships between phenotypes, highlighting the potential contribution of changes to genomic elements that contribute to the combinatorial regulation of genes that affect normal muscle function and may thus contribute to the severity of neuromuscular diseases[52–55].

### The pathways (and drugs) in common between GS and NMD share relationships underpinning the mechanisms of both age (sarcopenia) and disease (NMD) related muscle weakness

The mTOR signaling pathway (hsa04150), axon guidance (hsa04360) and alcoholism (hsa05034) pathways contain eGenes whose regulation is associated with genetic variants from each of the four phenotypes. The axon guidance pathway guides axon outgrowth and plays a pivotal role in forming a properly functional nervous system[56]. Axon guidance proteins involved in this pathway are responsible for neural circuit development and the changes in the expression and function of these proteins can cause neurological diseases. For example, increased expression of Sema3A and EPHA4 causes de-adhesion of neuromuscular junctions, leading to muscle denervation in ALS[57]. Changes in the mTOR signaling pathway (e.g. the loss of mTOR in skeletal muscle) or alterations to mTOR-mediated processes can contribute to myopathy and other skeletal muscle pathologies[58]. The mTOR protein coding gene expression correlates with the MS-associated variant rs3748817. Similarly, the overall function of the mTOR pathway is likely affected by alterations to RNF152, V-ATPase and WNT, which are eGenes for GS-, MG- and ALS-specific variants, respectively. Superficially the co-occurrence of eGenes within the KEGG-defined alcoholism pathway would appear to be inconsistent with our hypothesis. However, this pathway involves dopamine release, PKA signalling and activation of CREB-mediated gene expression. CREB activation promotes regeneration in instances of muscle damage[59]. Collectively, these findings are consistent with these pathways contributing core functions that mechanistically contribute to the maintenance and development of muscle strength.

We identified druggable eGenes and pathways for the MG, MS, ALS and GS phenotypes. Crucially, the overlapping druggable pathways are not simply the result of the presence of shared druggable eGenes. For instance, we identified co-occurring druggable eGenes (i.e. MG: *LTA;* and ALS: *CX3CR1*) within the cytokine-cytokine receptor interaction pathway. The co-occurrence of shared and phenotype-specific eGenes within these pathway overlaps informs on potential pharmacological side-effects associated with therapeutic treatment[60]. For example, vandetanib inhibits EPHA6 (receptor protein kinase) in the axon signaling pathway. However, in the presence of genetic variants that affect other genes within the axon signalling pathway (e.g. MS-associated eGene ERK), EPHA6 disruption could propagate through the axon signalling pathway to trigger the risk pathway for MS.

### High resolution maps of Hi-C connections in Immortalised Cell Lines overlap many, but not all, psoas Muscle-specific Hi-C connections

Overall, the SNP-gene pairs were observed across most of the Hi-C datasets used in this study. This is likely due to the conservation of topologically associated domains (TADs) across cell types and lineages[61,62]. However, a subset of SNP-gene interactions was only captured in the psoas muscle Hi-C data set, suggesting that gene regulatory interactions may be tissue specific in nature. Crucially, the biological pathways that contained co-occurring eGenes, which were specific to psoas muscle, represented a subset of the pathways identified from the immortalized cell line Hi-C datasets. Notably, none of the biological pathways containing co-occurring eGenes, identified from the psoas muscle Hi-C dataset, contained shared eGenes. For example, the Ras signaling pathway (required for the differentiation of slow muscle fibres by inducing slow motor neurons)[63] contains specific eGenes associated with: GS - *TGFA, RASGRF1*; MS- *RASGRP1, MAPK3, MAPK1, NFKB1, PLCG1, FLT1*; and ALS- *MRAS, TIAM1, FGF12*. Collectively, we contend that these observations reflect the tissue-specific impacts of the genetic variants and functional regulatory networks.

There are alternative explanations for the identification of more eGenes using Hi-C data from the immortalized cell lines[41] than from the psoas muscle[42]. The simplest explanation is that the chromatin interaction patterns emerge from the underlying nuclear functions and thus represent the different transcriptional networks in the pluripotent and differentiated cells[64]. Secondly, there is a strong linear relationship between the number of samples in GTEx and the number of eQTLs that are identified. Once saturated this relationship should reach an asymptote. Therefore, it remains possible that the tissue-specific pattern we are observing for the eGenes is due to under-sampling. Thirdly, cell-type-specific patterns of gene expression[65,66] may be lost due to tissue averaging. Fourthly, variation in the experimental Hi-C protocols used to capture the genome organization[41,42] may have resulted in false negatives in the psoas muscle data set. Despite these limitations, the co-occurrence of eGenes within biological pathways – in the presence of limited numbers of shared eGenes – indicates the utility of this approach for the identification of pathways underlying phenotype-specific characteristics.

## Conclusion

In conclusion, we have identified the existence of common genetic mechanisms by which neuromuscular diseases and aging cause muscle weakness and wasting. The shared gene regulatory mechanisms and the shared pathways identified between the phenotypes represent novel therapeutic targets and highlight possible mechanisms for side-effects and other complications.

## Code availability

CoDeS3D pipeline is available at https://github.com/Genome3d/codes3d-v1. R and python scripts used to clean the data and generate the figures are available in figshare with the identifier https://doi.org/10.17608/k6.auckland.9795350 R Studio version 1.2.1335 was used for all R scripts. Python version 2.7.15 was used for all the python scripts.

## Data availability

Supplementary Data 1 is available in figshare with the identifier https://doi.org/10.17608/k6.auckland.9789269. Supplementary Data 2 is available in figshare with the identifier https://doi.org/10.17608/k6.auckland.9789299. Supplementary Data 3 is available in figshare with the identifier https://doi.org/10.17608/k6.auckland.9789314. Supplementary Data 4 is available in figshare with the identifier https://doi.org/10.17608/k6.auckland.9789416. Supplementary Data 5 is available in figshare with the identifier https://doi.org/10.17608/k6.auckland.9789425. Supplementary Data 6 is available in figshare with the identifier https://doi.org/10.17608/k6.auckland.9789440. Supplementary Data 7 is available in figshare with the identifier https://doi.org/10.17608/k6.auckland.9789446. Supplementary Data 8 is available in figshare with the identifier https://doi.org/10.17608/k6.auckland.9789458. Supplementary Data 9 is available in figshare with the identifier https://doi.org/10.17608/k6.auckland.9789467. Supplementary Data 10 is available in figshare with the identifier https://doi.org/10.17608/k6.auckland.9790688. Supplementary Data 11 is available in figshare with the identifier https://doi.org/10.17608/k6.auckland.9790709. Human genome build hg19 (GRChr37) was downloaded from http://ftp.ensembl.org/pub/release-75/fasta/homo_sapiens/. SNP annotations (human genome, build hg19) were obtained from https://ftp.ncbi.nih.gov/snp/organisms/human_9606_b151_GRCh37p13/. Gene annotations were downloaded from http://www.gtexportal.org/static/datasets/gtex_analysis_v7/reference/gencode.v19.genes.v7.patched_contigs.gtf.gz.

## Supplementary figures

**Supplementary Figure 1.**
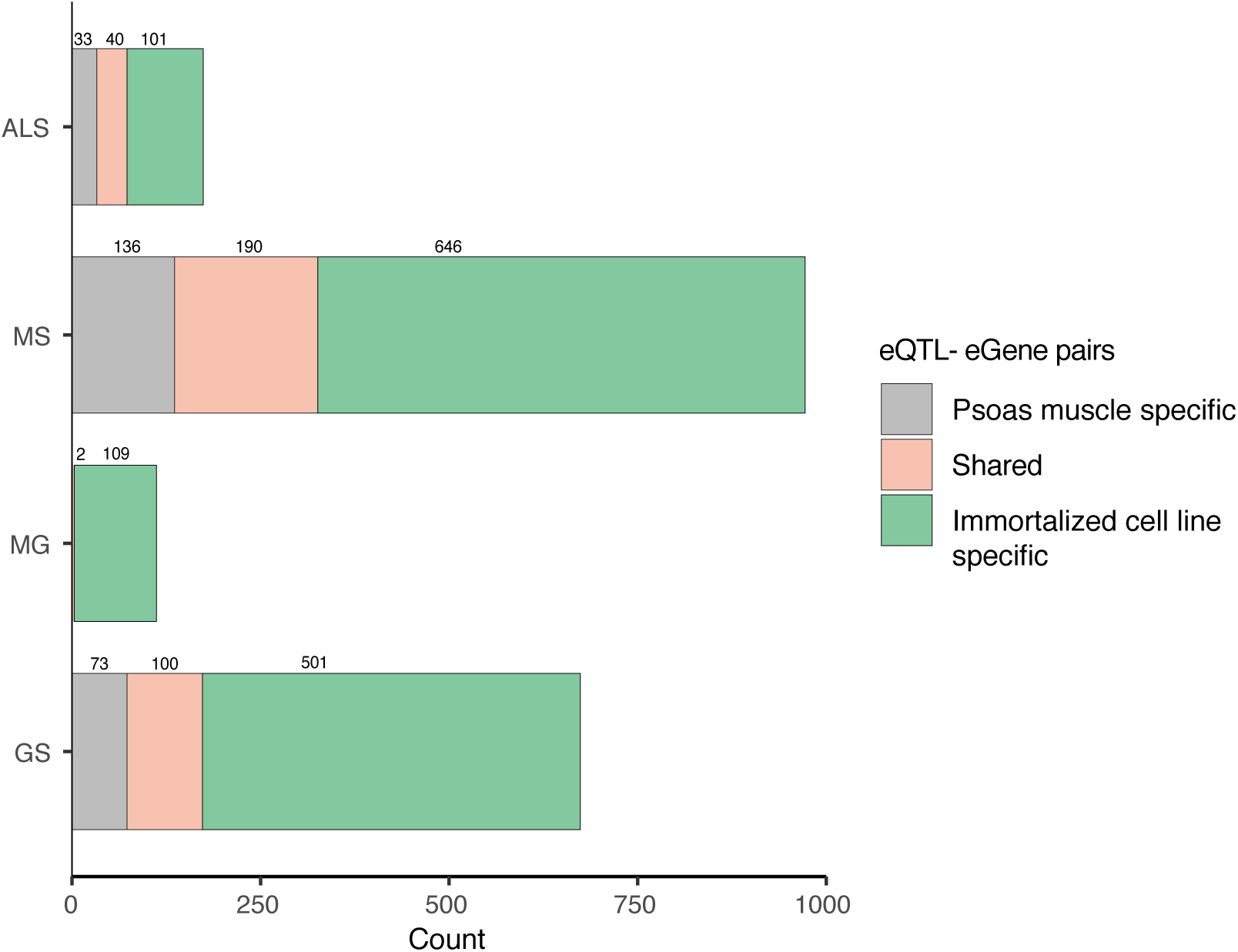
Considerable number of eQTL-eGene interactions are unique to psoas muscle Hi-C dataset. In all four phenotypes studied, more eQTL-eGene interactions were identified in the immortalized cell line Hi-C dataset, while the small number of interactions identified in the psoas muscle Hi-C dataset might represent tissue-specific chromatin interactions.

**Supplementary Figure 2.**
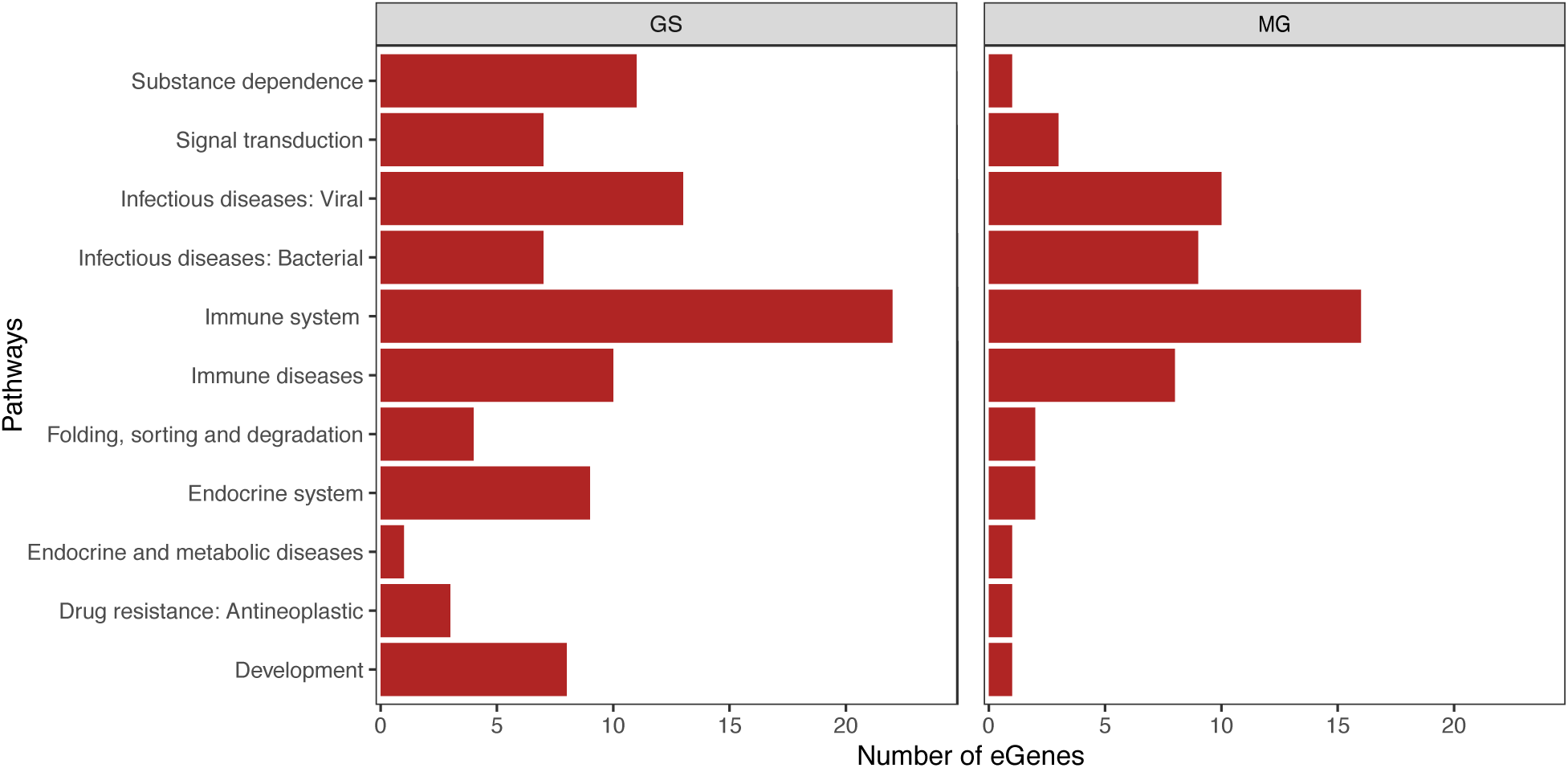
GS and MG eGenes identified from immortalized cell lines share pathways. Twenty four pathways **(Supplementary Table 7a)** classified under these 11 categories (y-axis) were shared by GS and MG eGenes. Red bars indicate the number of eGenes participating in those pathways from GS (left) and MG (right).

**Supplementary Figure 3.**
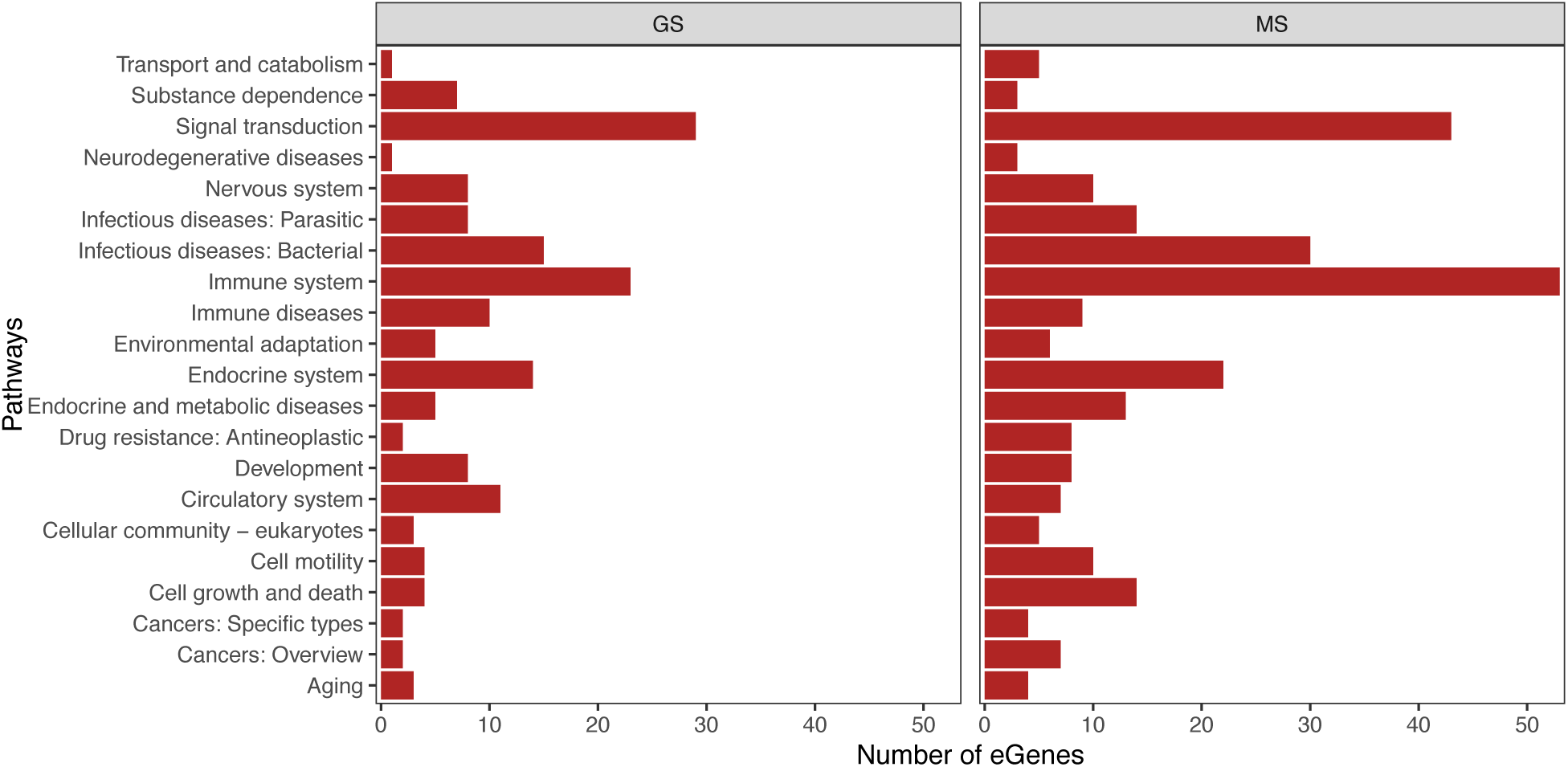
GS and MS eGenes identified from immortalized cell line share pathways. Sixty nine pathways **(Supplementary Table 7b)** classified under these 21 categories (y-axis) were found to be shared by GS and MG eGenes. Red bars indicate the number of eGenes participating in those pathways from GS (left) and MS (right).

**Supplementary Figure 4.**
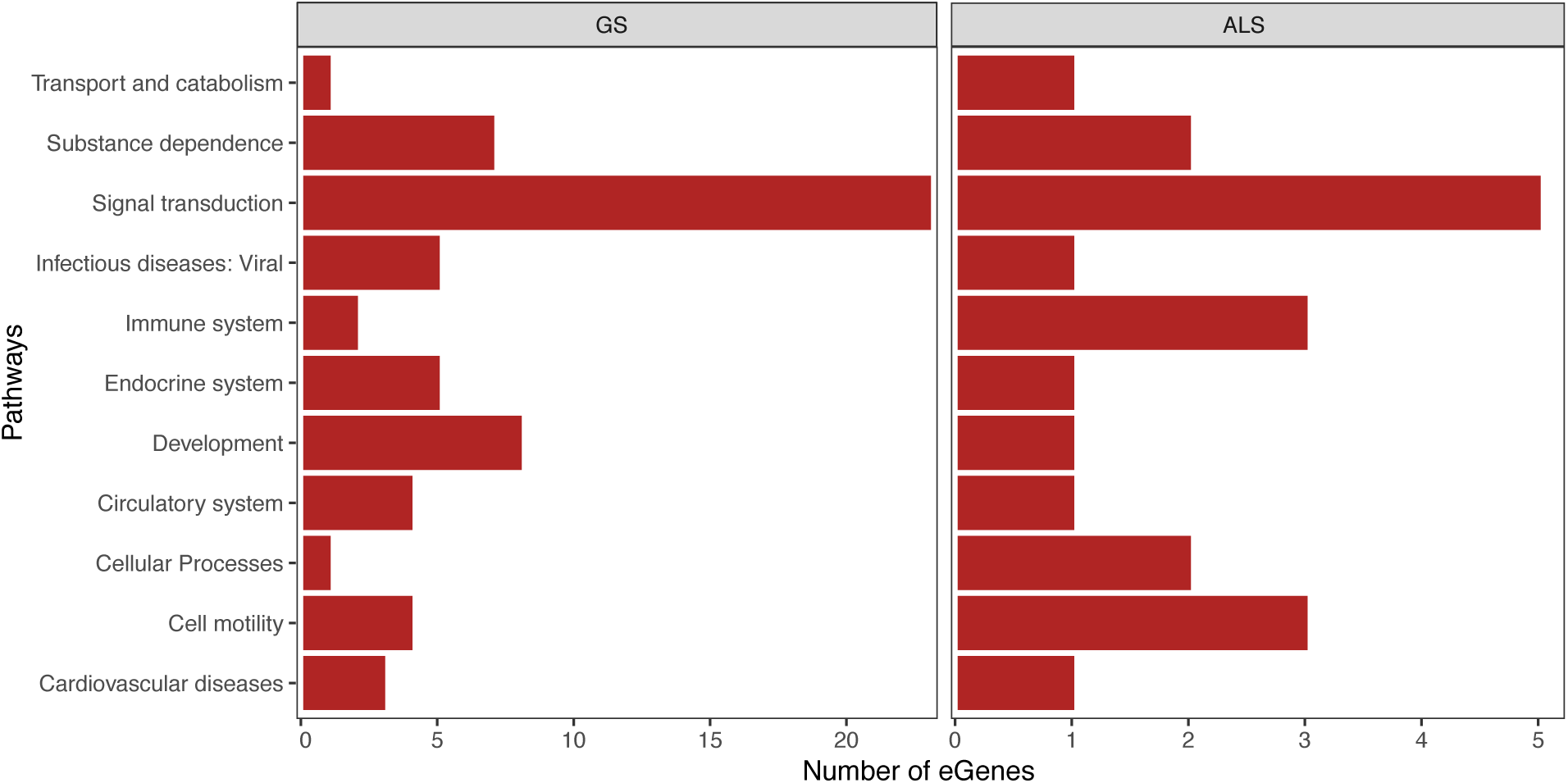
GS and ALS eGenes identified from immortalized cell lines share pathways. Seventeen pathways **(Supplementary Table 7c)** classified under these 11 categories (y-axis) were shared by GS and ALS eGenes. Red bars indicate the number of eGenes participating in those pathways from GS (left) and ALS (right).

**Supplementary Figure 5.**
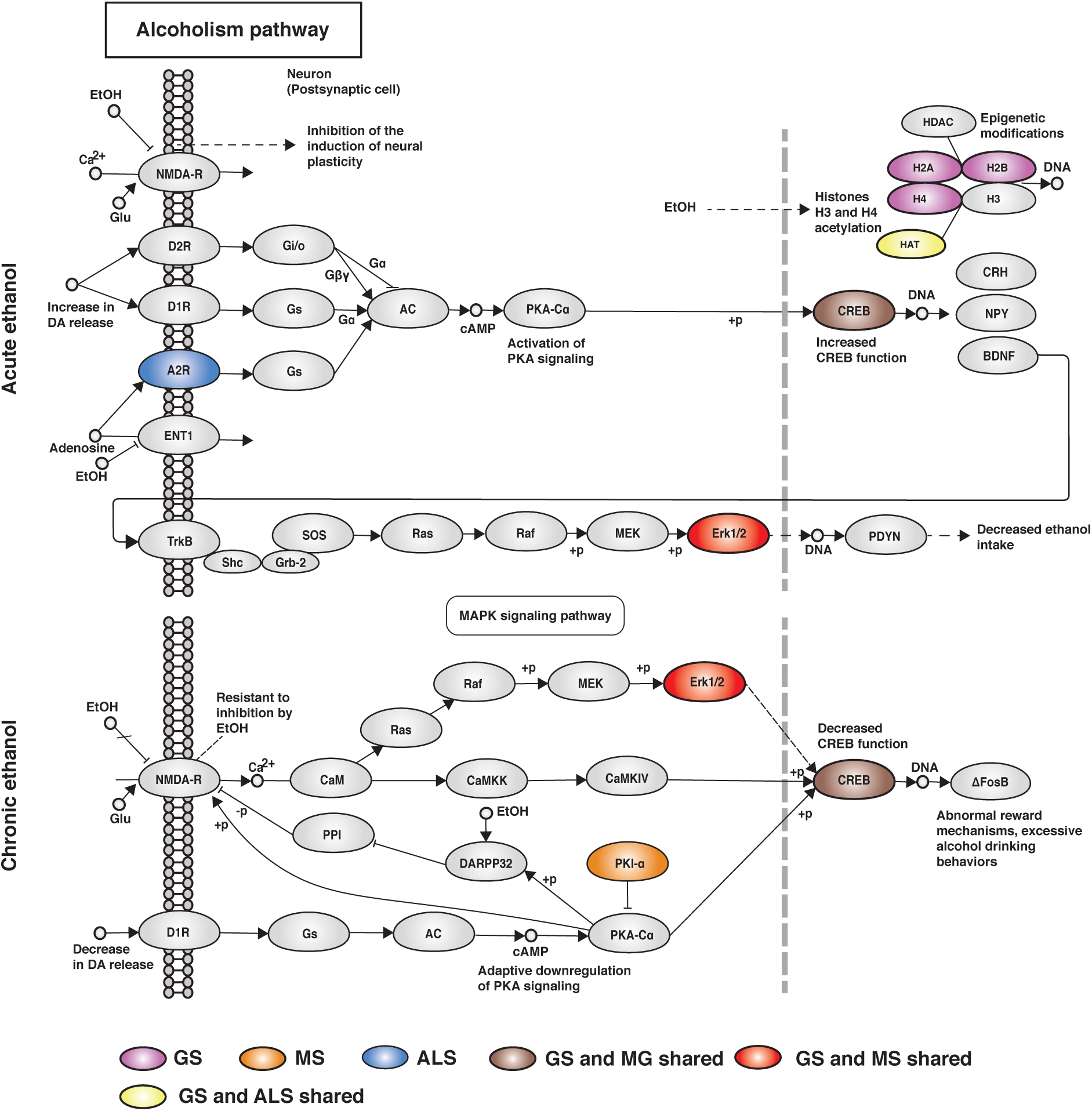
eGenes from all four phenotypes (GS, MG, MS and ALS) co-occurred within the alcoholism pathway. Chronic exposure to alcohol leads to pathological changes in several tissues and organs. Alcohol primarily impairs global protein synthesis under basal conditions as well as in response to the anabolic stimuli including muscle contraction^58^. GS-, MG-, MS- and ALS-associated eQTLs spatially regulate several eGenes that encode proteins involved in both acute and chronic alcohol-mediated signaling.

## Acknowledgments

The authors would like to thank the Genomics and Systems Biology Group (Liggins Institute) for useful discussions. This study was funded by a Ministry of Business, Innovation and Employment Catalyst grant (The New Zealand-Australia LifeCourse Collaboration on Genes, Environment, Nutrition and Obesity; UOAX1611; to JOS and SG. JOS and WS are funded by a Royal Society of New Zealand Marsden Fund [Grant 16-UOO-072]. We would like to thank Kane Hadley and Jared Nedzel from Genotype-Tissue Expression (GTE×) consortium for their technical support and the funders of GTEx Project - common Fund of the Office of the Director of the National Institutes of Health, and by National Cancer Institute, National Human Genome Research Institute, National Heart, Lung, and Blood Institute, National Institute on Drug Abuse, National Institute of Mental Health, National Institute of Neurological Disorders and Stroke.

## Author contributions

S.G performed analysis, interpreted data and wrote the manuscript. W.S contributed to data interpretation and commented on the manuscript. M.W commented on the manuscript. D.C.S commented on the manuscript. E.L.S contributed to data interpretation and commented on the manuscript. J.M.O directed the study, co-wrote the manuscript, contributed to data interpretation and commented on the manuscript. JOS is guarantor for this article.

